# Functional characterisation of the Nep1-like protein effectors of the necrotrophic pathogen - *Alternaria brassicae*

**DOI:** 10.1101/2021.07.04.450447

**Authors:** Deepak Duhan, Shivani Gajbhiye, Rajdeep Jaswal, Ravindra Pal Singh, Tilak Raj Sharma, Sivasubramanian Rajarammohan

## Abstract

*Alternaria brassicae* is an important necrotrophic pathogen that infects the *Brassicaceae* family. *Alternaria brassicae*, like other necrotrophs, also secretes various proteinaceous effectors and metabolites that cause cell death to establish itself in the host. However, there has been no systematic study of *A. brassicae* effectors and their roles in pathogenesis. The availability of the genome sequence of *A. brassicae* in public domain has enabled the search for effectors and their functional characterisation. Nep1-like proteins are a superfamily of proteins that induce necrosis and ethylene biosynthesis. They have been reported from a variety of microbes including bacteria, fungi, and oomycetes. In this study, we identified two NLPs from *A. brassicae* viz. AbrNLP1 and AbrNLP2 and functionally characterised them. Although both AbrNLPs were found to be secretory in nature, they localised differentially inside the plant. AbrNLP2 was found to induce necrosis in both host and nonhost species, while AbrNLP1 could not induce necrosis in both species. Additionally, AbrNLP2 was shown to induce pathogen-associated molecular pattern (PAMP)-triggered immunity in both host and nonhost species. Overall, our study indicates that AbrNLPs are functionally and spatially (subcellular location) distinct and may play different but important roles during the pathogenesis of *A. brassicae*.

## Introduction

Plant pathogens secrete various proteins, secondary metabolites, and other small molecules to colonize the host by suppressing host defences [1, 2]. In contrast, many pathogens also produce proteins that trigger cell death/necrosis and associated responses in the host. Various filamentous fungal and oomycetes pathogens can induce cell death in the plants through the secretion of cell death inducing proteins/effectors [3-5]. Cell death (either necrotic or apoptotic-like) plays a central role in plant-pathogen interactions either by restricting pathogen growth (in case of biotrophs) or by providing a nutrient source for the proliferation of the pathogen (in case of necrotrophs). Many well-known necrotrophic pathogens such as *Botrytis cinerea, Sclerotinia sclerotiorum, Alternaria brassicae* have been shown to induce cell death in the host and effective induction of cell death leads to susceptibility [6-8].

The genus of *Alternaria* contains many important plant pathogens causing diseases in agronomically important cereal, vegetable, oilseed, and fruit crops. These species are known to be one of the major producers of host-specific toxins and metabolites that cause cell death and help in disease progression. *Alternaria* spp. produce chemically diverse secondary metabolites that range from low molecular weight secondary metabolites to peptides. These toxins can be host-specific as well as non-specific. Tenuazonic acid, tentoxin, and brefeldin A are some examples of non-specific toxins secreted by *Alternaria* spp., which are phytotoxic in nature [9, 10]. Host-specific toxins (HSTs) such as AK-toxin, AF-toxin, ACT-toxin, AM-toxin, AAL-toxin are secreted by various pathotypes of *A. alternata* depending on the host species they infect [11-13]. Although many studies have shown the role of these secondary metabolite toxins in pathogenesis of *Alternaria* spp., very little is known about the molecular mechanisms underlying pathogenesis of other *Alternaria* species such as *Alternaria brassicae* and *Alternaria brassicicola*. Destruxin B was considered to be an HST of *A. brassicae*, but later studies proved that destruxin B only aids in the pathogenesis of *A. brassicae* [14]. Recent studies have revealed that necrotrophic pathogens also secrete small proteinaceous effectors that can function as virulence factors. Some prominent examples of proteinaceous necrotrophic effectors that induce cell death include PtrToxA, SnToxA, BcCFEM1, BcXYG1, and the NEP-like proteins in various necrotrophs [15-17].

Nep1-like proteins (NLPs) are a group of proteins identifiable by the presence of a common necrosis-inducing NPP1 domain [18, 19]. NLPs constitute a superfamily of proteins, which are present in both prokaryotes and eukaryotes [3, 20, 21]. The founding member of this family is a 24-kDa protein (Nep1) that was identified in *Fusarium oxysporum* that could cause necrosis and induce ethylene biosynthesis in dicots [22]. The NPP1 domain present in all NLPs is characterised by a conserved heptapeptide motif – GHRHDWE, and N-terminal conserved cysteine residues. Based on the number of cysteine residues, NLPs are classified as type I (two conserved cysteine residues) or type II (four conserved cysteine residues) [3]. Most NLPs identified in oomycetes and fungi trigger cell death and also act as pathogen-associated molecular patterns (PAMPs), thereby activating PAMP-triggered immunity (PTI) [23-25]. However, the role of NLPs in virulence is conflicting with NLPs being dispensable for virulence in *Zymoseptoria tritici* and *B. cinerea* and known to accelerate disease and pathogen growth in *Colletotrichum coccodes, Phytophthora capsici* and *Pythium* species [26-28]. However, there have been no studies on the identification or functional characterisation of NLPs from the genus of *Alternaria*.

The current study was undertaken to identify and functionally characterise the NLPs in *A. brassicae*, a notorious necrotrophic pathogen of the *Brassica* crops. Specifically, we aimed to 1) identify the NLPs in *A. brassicae* and their phylogenetic relationship with other known NLPs, 2) to study the temporal expression pattern in the natural host - *Brassica juncea*, 3) to determine their ability to cause necrosis in host as well as nonhost species, 4) to identify their subcellular localisation, and 5) to analyse if the NLPs in *A. brassicae* also acted as PAMPs and induced PTI in the host.

## Experimental Procedures

### Fungal strains, plants, and culture conditions

*A. brassicae* J3 strain was used for the infection assays in the expression analysis. The strain was grown on potato dextrose agar (PDA) plates at 22°C. For the transient expression assays, *N. benthamiana* wild type plants grown at 25 °C with photoperiod of 10h Light/14h dark were used. Similarly, *B. juncea* var. Varuna was grown at 25 °C with photoperiod of 10h Light/14h dark and used for the infection and infiltration assays.

### Identification and *in silico* analysis of *A. brassicae* NLPs

NLP genes in *A. brassicae* were identified by a BLAST search of the *A. brassicae* proteome using the NPP1 domain (Pfam: PF05630) as a query. Signal peptides were predicted using SignalP 5 [29]. Phylogenetic analysis was performed using MEGA X [30]. Protein sequences of other *Ascomycetes* members were retrieved using a similar BLAST search as described above against the nr database (Supplementary Table S1). Protein sequences were aligned using MUSCLE and outliers with too many gaps were removed from the analysis. The WAG+G substitution model was selected since it had the least Bayesian Information Criterion (BIC) value among the 56 amino acid substitution models tested. A phylogenetic tree was constructed using the WAG+G model in MEGA X and the final tree was visualized in iTOL [31].

### Gene expression analysis

Fungal cultures were established and plant infection assays were carried out as described earlier [32]. Leaves were collected at two and four days post infection (2 & 4 dpi). Fifteen-day old *A. brassicae* mycelia growing on PDA was also harvested (*in vitro* sample). Total RNA was extracted from 2-3 leaves collected from three individual plants in each experiment using the RNeasy plant mini kit according to the manufacturer’s recommendation (Qiagen, Gaithersburg, MD, U.S.A.). First-strand cDNA was synthesised from 1 μg of total RNA using PrimeScript 1^st^-Strand cDNA Synthesis Kit (Takara Bio, Japan) as per manufacturer’s protocol. PCRs were carried out using the standard setting in a CFX Connect Real-Time PCR System (BIO-RAD, California, USA). The housekeeping gene AbrActin was used as the endogenous control. Further, to calculate the relative expression levels, the dCt values of the infected samples were normalised to the *in vitro* sample (*A. brassicae* mycelia on PDA). Relative expression values were calculated using the delta delta Ct method [33]. All expression experiments were carried out in three biological replicates. Every biological replicate also consisted of three technical replicates. Sequences of primers used in qPCR have been listed in Supplementary Table S2.

### Yeast secretion trap (YST) assay

Experimental validation of the secretory nature of the AbrNLPs was done using yeast secretion trap assay. The *AbrNLP* genes were cloned into the pSUC-GW vector. Yeast strain YTK12, which is SUC2 negative, was transformed using the Fast Yeast Transformation kit (G-Biosciences, MO, USA). All transformants were selected on a yeast minimal medium without tryptophan. To check for secretion, 20 μl overnight yeast cultures with A600 ∼ 1.0 was plated onto YPSA medium (1% yeast extract, 2% peptone, 2% sucrose and 1 μg/ml antimycin A). The untransformed YTK12 strain, which is unable to grow on a sucrose medium, was used as a negative control.

### Transient expression of AbrNLP1 and AbrNLP2

#### Molecular cloning and sequencing

The *AbrNLP* genes (with signal peptide and without stop codon) were amplified from cDNA of *A. brassicae* infected leaves of *B. juncea* and cloned in the pENTR entry vector. The entry clone was then Sanger sequenced to confirm the cDNA sequences of *AbrNLPs*. The entry clones were then transferred to different destination vectors using LR clonase II (Gateway cloning technology) for functional assays. Three destination vectors were used viz. pGWB408, pGWB441, and pDEST17. Positive clones of the destination vectors were confirmed using colony PCR and restriction digestion. The confirmed vectors were then mobilized into the *Agrobacterium tumefaciens* GV3101 strain.

#### Cell death assays

Primary cultures of pGWB408-AbrNLP1 and pGWB408-AbrNLP2 grown overnight were used for inoculating secondary cultures, which were grown till they reached an OD_600_ of 0.8-1. The pellets from the secondary cultures were resuspended in the resuspension solution (MES– 10 mM, MgCl_2_ – 10 mM, and Acetosyringone – 200 μM). The resuspended cultures were incubated at room temperature for 3-4 hours prior to infiltration in 5-6 weeks old *Nicotiana benthamiana* and *B. juncea* leaves. Cell death was observed 3-4 days post infiltration.

#### Subcellular localisation

The AbrNLPs were cloned into the pGWB441 vector to have a C-terminal yellow fluorescent protein (YFP) fusion using Gateway Cloning. Primary cultures of pGWB441-AbrNLP1 and pGWB441-AbrNLP2 grown overnight were used for inoculating secondary cultures, which were grown till they reached an OD_600_ of 0.8-1. The pellets from the secondary cultures were resuspended in the resuspension solution (MES – 10 mM, MgCl2 – 10 mM, and Acetosyringone – 200 μM). The resuspended cultures were incubated at room temperature for 3-4 hours prior to infiltration in 5-6 weeks old *N. benthamiana* leaves. The subcellular localisation of YFP-tagged proteins was examined 2-3 days post infiltration using a Carl Zeiss confocal microscope (LSM880).

#### Heterologous expression and purification

For protein expression, the entry clone was mobilized into the pDEST17 destination vector. Positive clones were transformed into BL21(DE3)pLysS competent cells. The cultures were induced with 0.4 mM IPTG at 22 °C. Following expression, the cells were harvested and cell lysis was carried out by chemical, enzymatic and mechanical treatment. For chemical lysis, cells were suspended in a lysis buffer (containing 100 mM NaCl, 50 mM Tris buffer of pH8.0, 200 μl Barry’s buffer and 2 mM PMSF). The cell suspensions were then subjected to lysis with 200 μg/gm lysozyme for 30 min followed by treatment with DNase I for 1 h at room temperature. The lysates were then sonicated with 40% amplitude, 10s ON/30s OFF cycle for 3 min on ice, followed by centrifugation at 10,000x g for 40 min at 4 °C. The supernatant was then subject to Ni-NTA purification using Ni Sepharose High-Performance resin (Cytiva Lifesciences, Marlborough, USA). The eluted purified protein was desalted to remove excess imidazole. The purified protein was further used for cytotoxicity and secreted peroxidase assays.

#### Cytotoxicity of AbrNLP2 and secreted peroxidase (sPOX) assay

Purified AbrNLP2 was directly infiltrated into the leaves of 5-6-week old *N. benthamiana* and *B. juncea* plants to check for cytotoxicity. The sPOX assay was carried out as described earlier with minor modifications [34]. Briefly, 5 mm leaf discs were excised from *N. benthamiana* and *B. juncea* plants and washed with distilled water for 2 hours with agitation in a 96 well plate. After washing, water was carefully removed out while minimizing the damage to the leaf discs. 100 µl of purified AbrNLP2 protein was added in the designated wells while remaining wells filled with buffer and incubated for 20 h with agitation. After incubation, the leaf discs were carefully removed from the wells and 50 µl of 5-aminosalicylic acid (1 mg/ml) supplemented with 0.01 % H_2_O_2_ was added in each well. After 3 minutes, 20 µl of 2N NaOH was added to each well to stop the reaction and absorbance was measured at OD_600_.

## Results

### The genome of *A. brassicae* encodes two NLPs

To identify NLPs in the genome of *A. brassicae*, we searched the whole protein dataset using the NPP1 domain as a query. We found two genes that contained the NPP1 domain viz. ABRSC02.1105 and ABRSC02.949 (hereafter referred to as AbrNLP1 and AbrNLP2 respectively). The two AbrNLPs identified were highly diverged sharing only ∼40% similarity (Figure 1A). Additionally, both the AbrNLPs contained the two conserved cysteine residues and the heptapeptide motif (GHRHDWE), which classified them as a type I NLP [3]. In order to reconstruct the relationship between AbrNLPs and the NLPs identified in other *Ascomycetes*, a phylogenetic analysis was carried out. As expected the AbrNLP1 clustered along with NLP1 homologs from other *Alternaria* species, while AbrNLP2 clustered together with the NLP2 homologs (Figure 1B). Moreover, we found no correlation between the presence of the NLP proteins and the lifestyle of the fungal pathogens.

**Figure 1:**
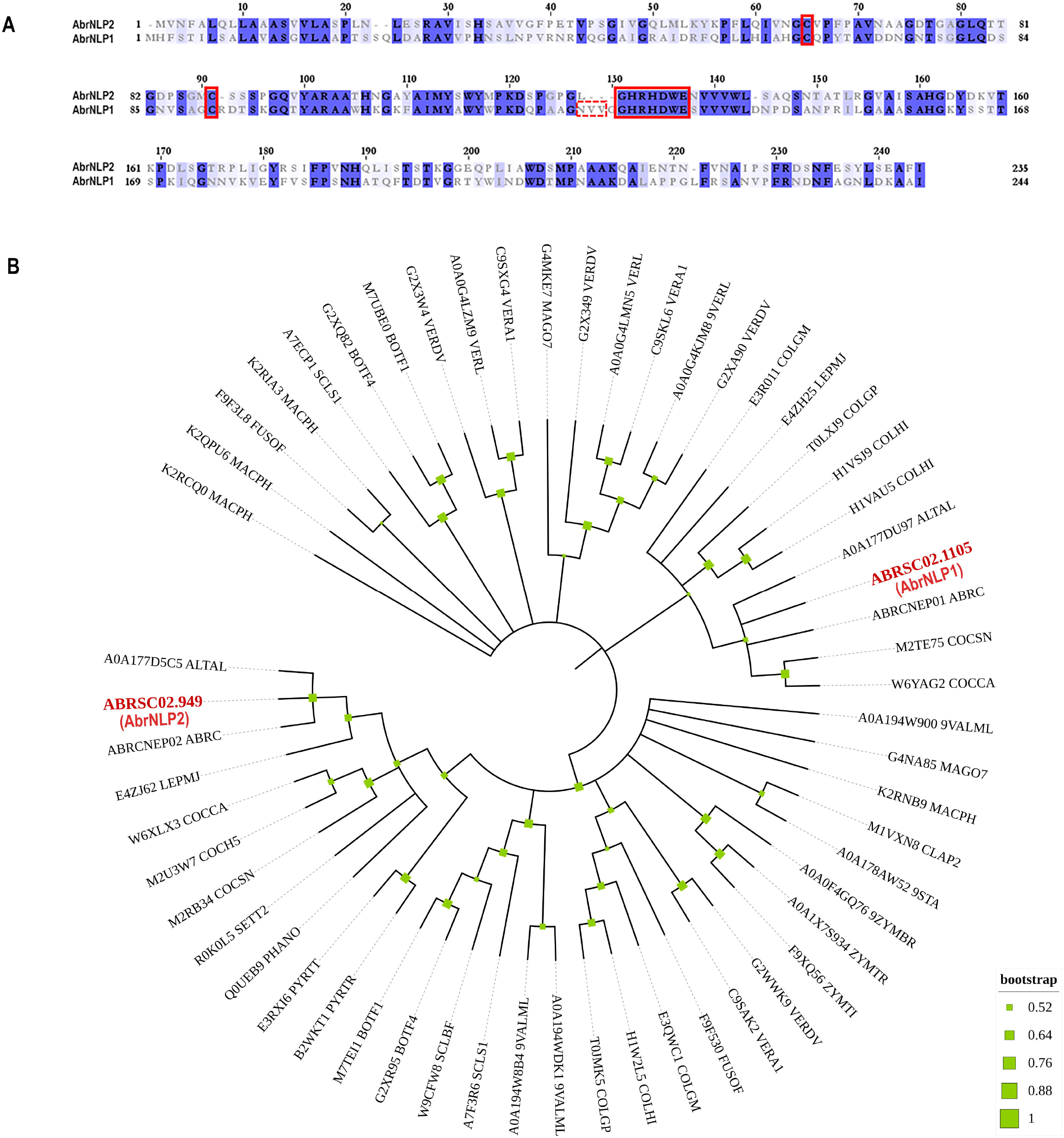
Sequence characteristics of AbrNLPs and their phylogenetic relationship. A) Multiple sequence alignment of NLPs from *A. brassicae*, B) Phylogenetic relationship of AbrNLP1 and AbrNLP2 compared to the NLPs of other pathogens was inferred by using the Maximum Likelihood method. The bootstrap consensus tree was inferred from 1000 replicates. Branches corresponding to partitions reproduced in less than 50% bootstrap replicates were collapsed. A discrete Gamma distribution was used to model evolutionary rate differences among sites (5 categories (+G, parameter = 1.4565)). All positions with less than 95% site coverage were eliminated, i.e., fewer than 5% alignment gaps, missing data, and ambiguous bases were allowed at any position.

### AbrNLP1 and AbrNLP2 both are induced upon infection in Arabidopsis & *B. juncea*

The expression of AbrNLP1 and AbrNLP2 was studied *in vitro* (on PDA plates) and over the course of infection in the natural host of *A. brassicae* i.e. *B. juncea*. Previous studies on the infection of *A. brassicae* have identified two critical time points viz. 2 and 4 days post infection [6], therefore we studied the expression of AbrNLP1 and AbrNLP2 at these time points. We found that both AbrNLP1 and AbrNLP2 are expressed *in vitro* i.e. in growing fungal mycelia on solid media. However, they are both upregulated during the infection process at 2 dpi and thereafter are downregulated at 4 dpi when infection is established (Figure 2).

**Figure 2:**
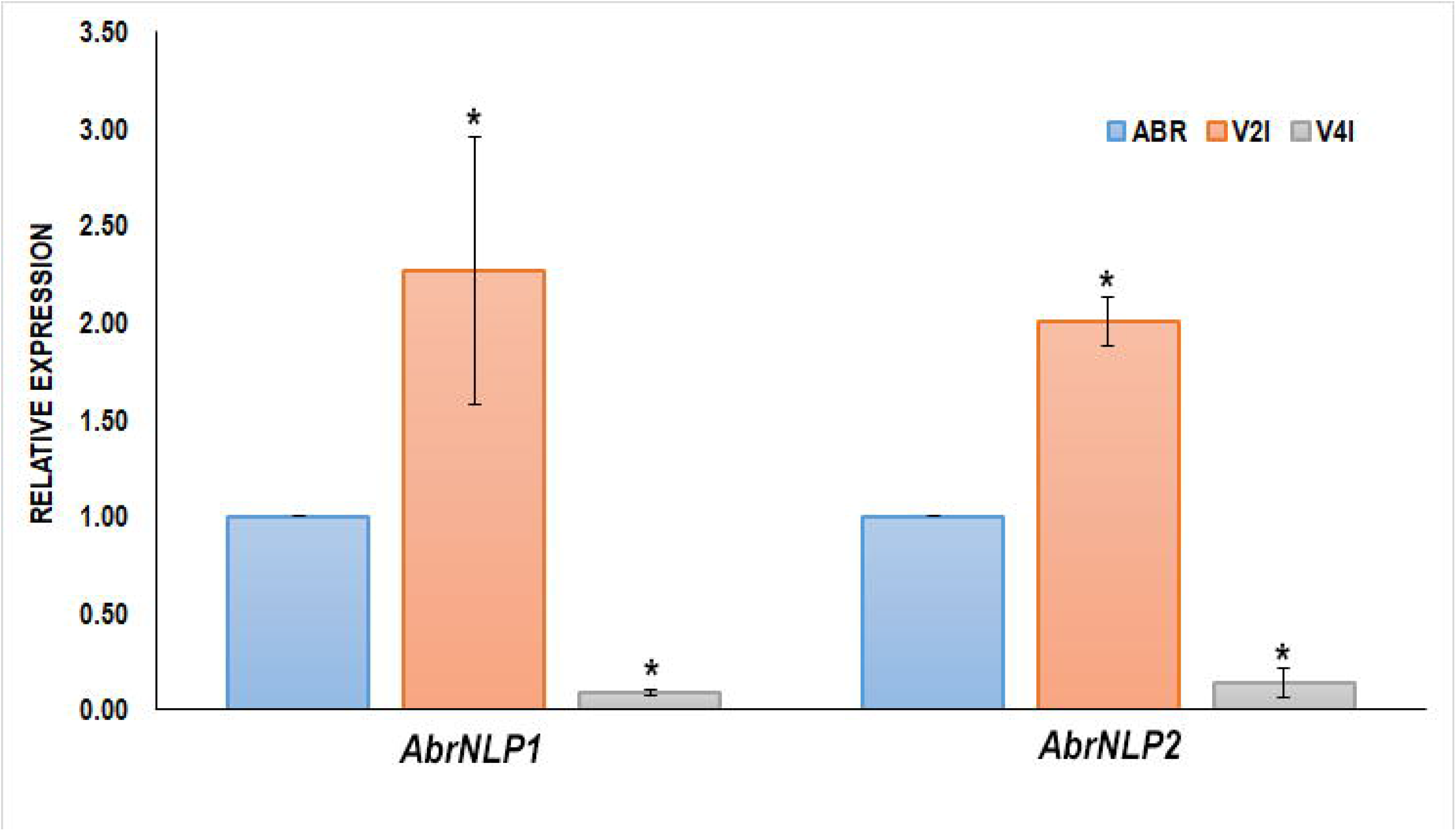
Expression of *AbrNLPs* during infection of *B. juncea*. Relative expression analysis of AbrNLP1 and AbrNLP2 in infected *B. juncea* (natural host) leaves 2 (V2I) and 4 (V4I) days post infection with respect to plate-grown fungi (ABR). AbrACTIN was used as an endogenous control. The mean values (± standard deviation) of three biological replicates are shown. *P < 0.05 by Mann–Whitney U-test.

### AbrNLP1 and AbrNLP2 contain bonafide signal peptides and are secreted outside the cell

Both the proteins, AbrNLP1 and AbrNLP2 were predicted to have an N-terminal signal peptide and most likely to be secreted. Further, AbrNLP1 and AbrNLP2 were predicted to be effectors by the EffectorP 2.0 tool [35]. In order to confirm the secretion of AbrNLP1 and AbrNLP2, a yeast invertase secretion assay was carried out. Both proteins grew on the sucrose selection media indicating that the N-terminal signal peptide resulted in the secretion of these proteins outside the cell (Figure 3).

**Figure 3:**
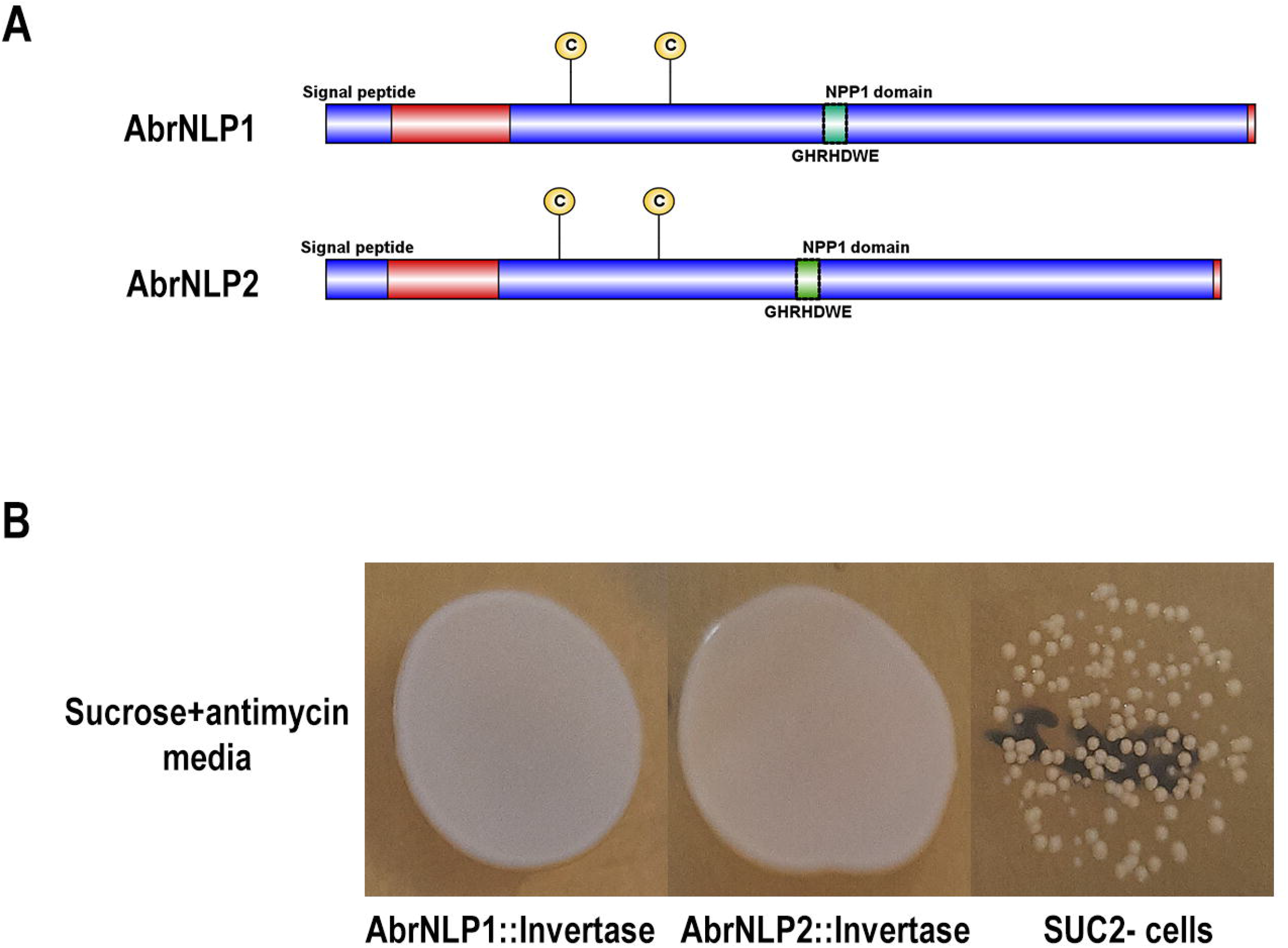
AbrNLP1 and AbrNLP2 are secreted proteins as determined by the YST assay. A) Schematic diagram of AbrNLP1 and AbrNLP2 indicating the signal peptides and NLP domains, B) Yeast strain SUC2^-^ was transformed with AbrNLP1::Invertase and AbrNLP2::Invertase, and grown on medium containing sucrose with or without antimycin. Untransformed SUC2^-^ was used as a negative control.

### AbrNLP2 induces cell death while AbrNLP1 induces only chlorosis in *N. benthamiana*, and *B. juncea*

To check AbrNLPs for their ability to induce necrosis, the full-length genes (along with their signal peptide) were transiently expressed in *N. benthamiana* and *B. juncea* (natural host). BAX (Bcl2-associated X), known to induce apoptotic-like cell death in plants, was used as a positive control. AbrNLP1 could not induce cell death in *N. benthamiana* or *B. juncea* even after 5-7 days of agroinfiltration. However, *N. benthamiana* leaves infiltrated with AbrNLP1 developed chlorosis after 3 days of infiltration, which was absent in the regions infiltrated with empty vector controls (Figure 4). AbrNLP2 induced cell death 2 days after agroinfiltration in *N. benthamiana* and *B. juncea*, although the extent of cell death was less compared to the positive control - BAX.

**Figure 4:**
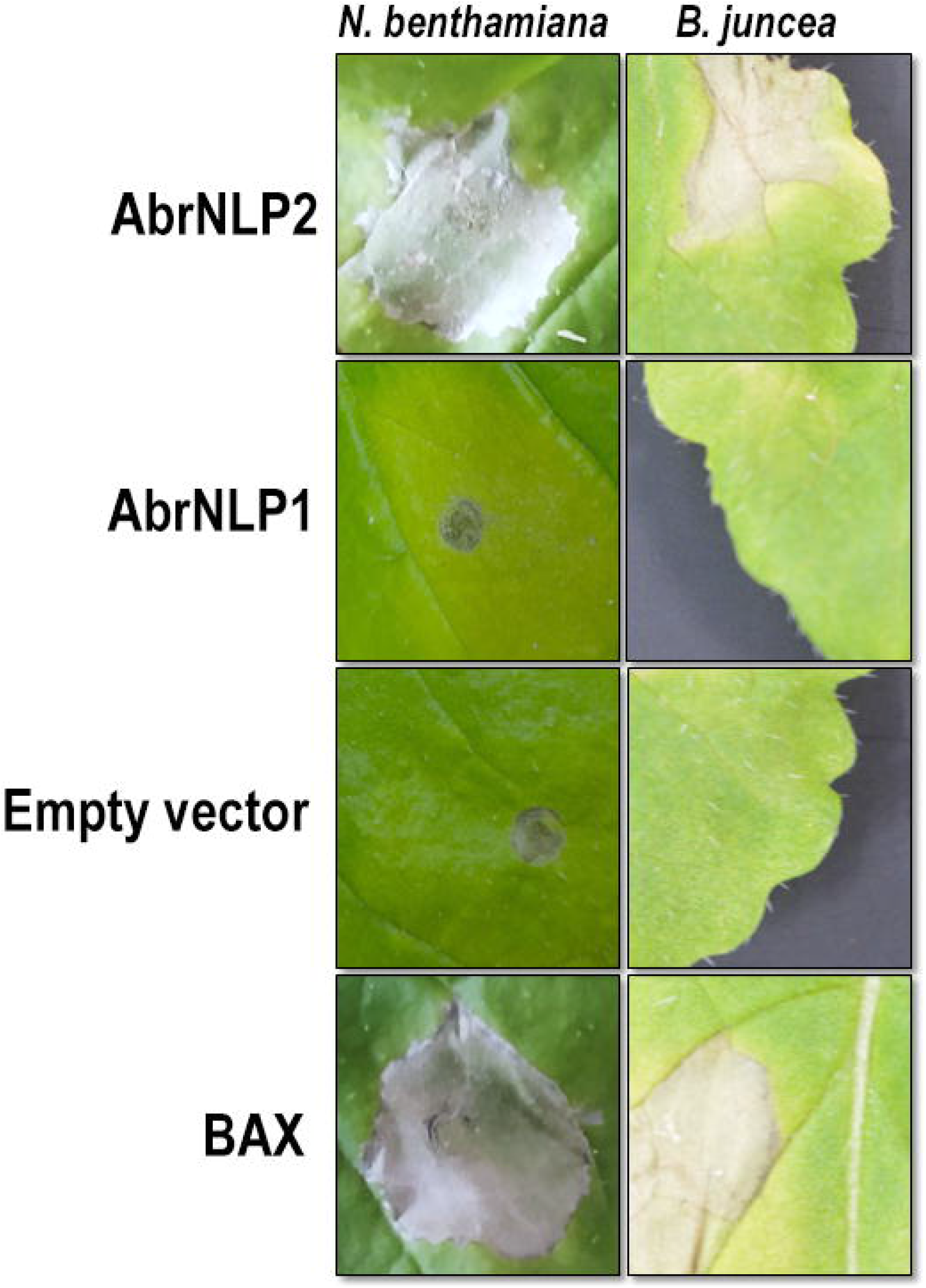
Agroinfiltration of AbrNLP1 and AbrNLP2 in *N. benthamiana* and *B. juncea*. Transient expression of AbrNLP1 and AbrNLP2 in both host and nonhost species. BAX was used as a positive control for induction of cell death and empty vector (pGWB408) was used as a negative control.

### Differential localisation of AbrNLP1 and AbrNLP2 in *N. benthamiana*

AbrNLP1 was predicted to be localised in the apoplast, while AbrNLP2 was predicted to be non-apoplastic by the ApoplastP tool [36]. In order to experimentally determine the subcellular localisation of AbrNLP1 and AbrNLP2, these proteins were transiently expressed in *N. benthamiana* with a C-terminal YFP tag. Full-length YFP under the 35S promoter was used as a positive control. AbrNLP1 was found to be localized in the plasma membrane and the apoplastic space between the cell junctions, whereas AbrNLP2 was localised to the plasma membrane, cytoplasm, and the nucleus as well (Figure 5) thereby confirming the predictions by the ApoplastP tool. However, AbrNLP2 does not contain any nuclear or organellar localisation sequences.

**Figure 5:**
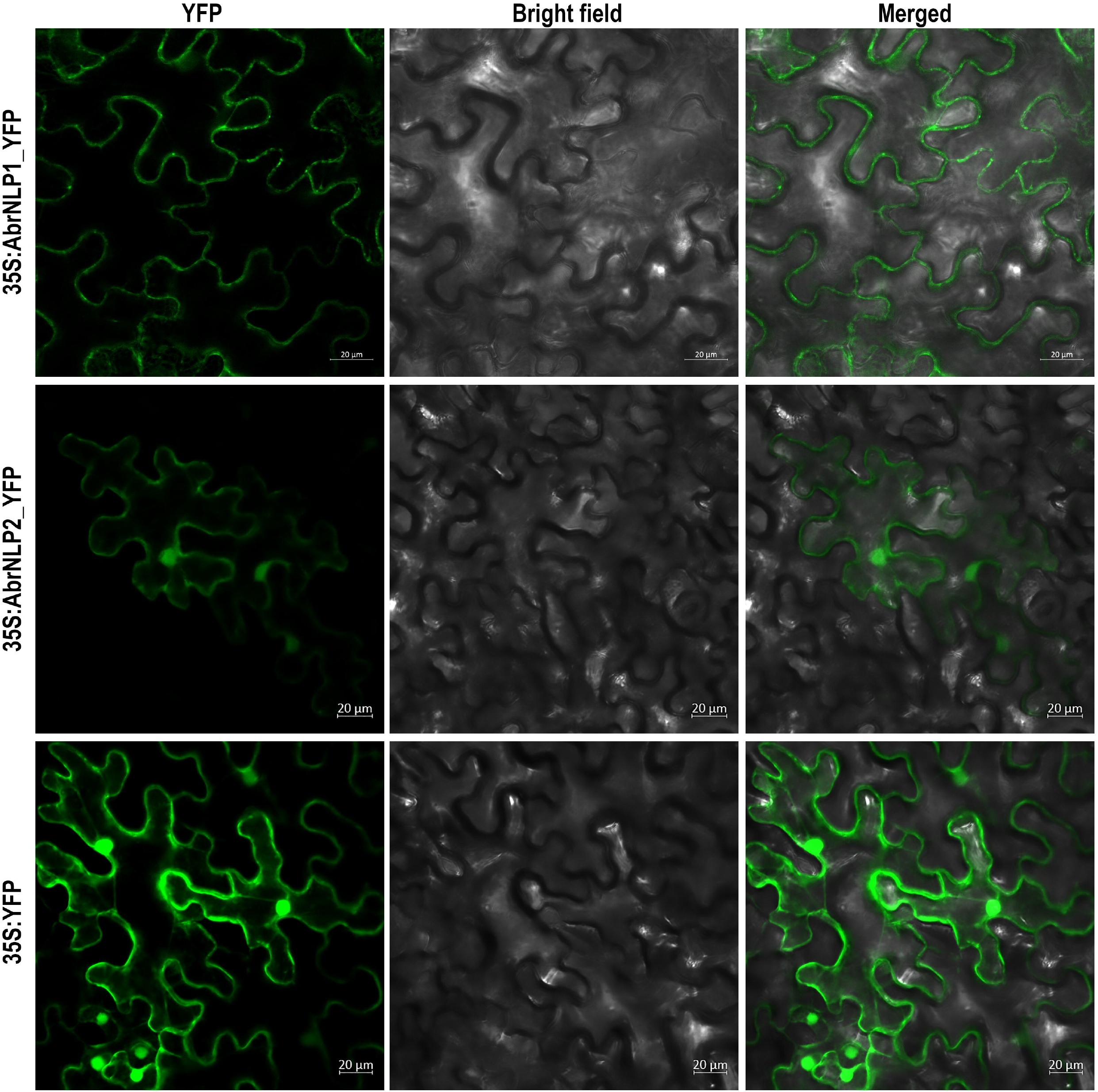
Subcellular localisation of YFP-tagged AbrNLP1 and AbrNLP2 in *N. benthamiana*. Confocal microscopy images of transiently expressed C-terminal YFP-tagged AbrNLPs in *N. benthamiana* 72 hpi. Full-length YFP was used as a positive control. scale bar = 20 µm.

### AbrNLP2 induces PTI responses in *N. benthamiana* and *B. juncea*

NLPs from various pathogens have been shown to be both cytotoxic as well as an inducer of PTI. Therefore, to check if AbrNLPs could also induce PTI responses in the host, we heterologously expressed AbrNLPs in *E. coli* and purified them (Figure 6A). However, AbrNLP1 was consistently found in the inclusion bodies in all the conditions and hence could not be purified. AbrNLP2 was purified and was used for both cytotoxic assays and a secreted peroxidase assay, which is an indicator of PTI response [34]. Purified AbrNLP2 protein could induce cell death in both *N. benthamiana* and *B. juncea* within 2 days of infiltration (Figure 6B). The same dilution was used for the sPOX assay, wherein AbrNLP2 treated leaf discs of *N. benthamiana* and *B. juncea* had significantly higher sPOX activity than the buffer treated leaf discs (Nb: p-value - 0.00587, t-stat - −3.38; Bj: p-value - 0.0103, t-stat - −2.97) indicating that AbrNLP2 is also capable of inducing PTI (Figure 6C).

**Figure 6:**
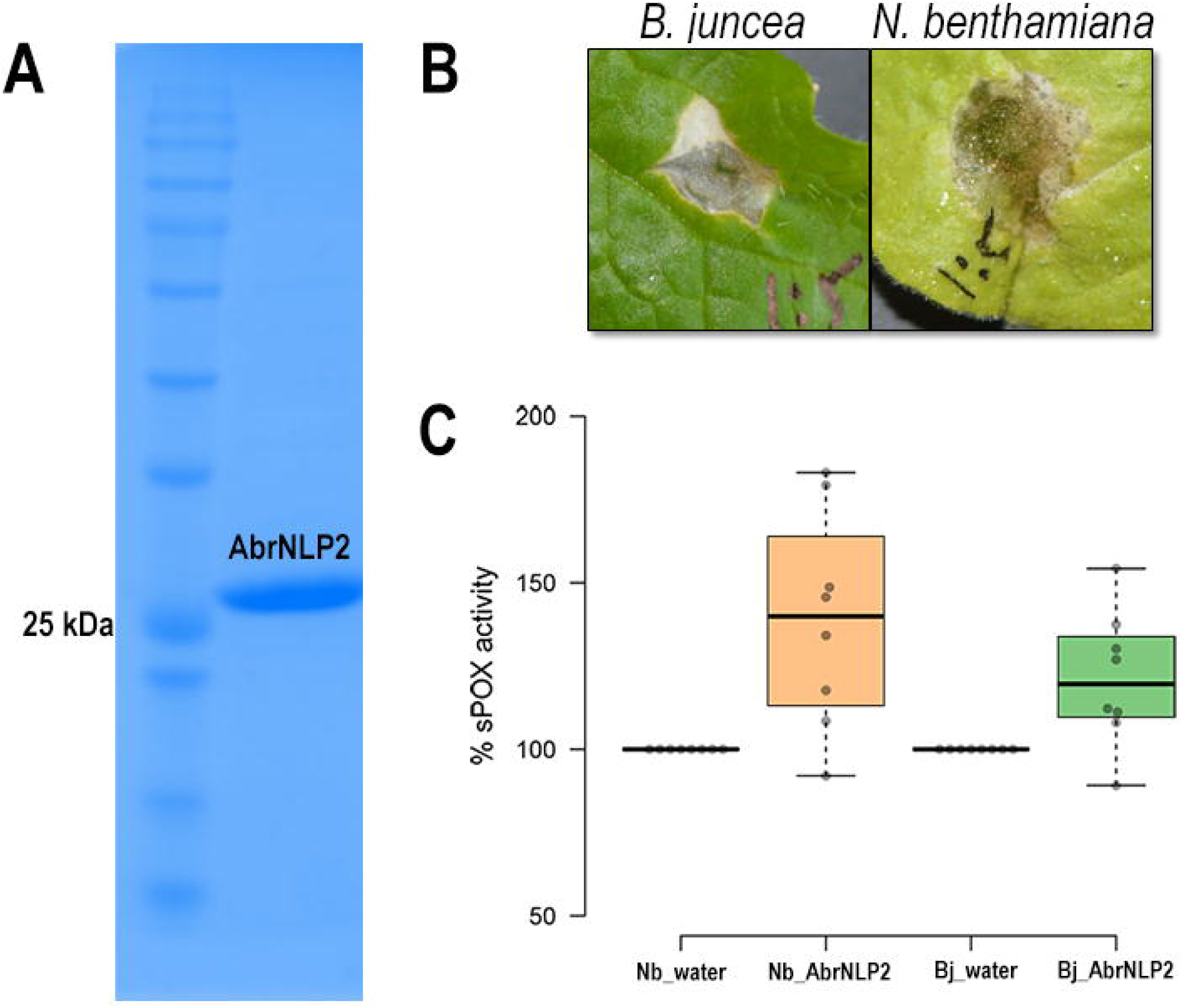
AbrNLP2 induces cell death and PTI responses. A) Ni-NTA based purification of AbrNLP2, B) Cell death induced by purified AbrNLP2 in *N. benthamiana* and *B. juncea*, C) %sPOX activity of *N. benthamiana* and *B. juncea* in response to purified AbrNLP2 as compared to water (control).

## Discussion

An emerging number of studies have shown that cell death-inducing proteins/effectors are required for pathogenicity, or contribute to the virulence of necrotrophic pathogens. Proteinaceous necrotrophic effectors have received less attention due to the gamut of earlier studies focussing on secondary metabolite toxins as the major pathogenicity factors in many of the necrotrophic pathogens [12, 13, 37]. Recent genome sequencing efforts and functional studies have shown that proteinaceous effectors of necrotrophic pathogens have a larger role to play in the pathogenicity than previously surmised [7, 38, 39].

NLPs are a superfamily of proteins found in pathogenic bacteria, fungi, and oomycetes and have been implicated in their virulence and disease pathogenesis. In the current work, we identified and characterised the NLPs in *A. brassicae*, an important pathogen belonging to the genus of *Alternaria*. We found that most species contained only 2 copies of the NLP genes within the genus of *Alternaria*, while some endophytes have only one (Supplementary Table S3). This is in contrast with oomycetes pathogens, which have an expanded repertoire of NLPs reaching up to 33 copies in *P. sojae* [40]. However, most broad range necrotrophs such as *B. cinerea, S. sclerotiorum*, and *A. brassicae* possess only two copies of NLPs.

Both the NLPs of *A. brassicae* were upregulated in the initial stages of infection, suggesting that AbrNLP1 and AbrNLP2 may have a role in pathogenesis. However, AbrNLP1 did not induce cell death *in N. benthamiana* or *B. juncea*. Most NLPs that have been identified in necrotrophs have a strong ability to induce necrosis. However, many non-cytotoxic NLPs have been found in hemibiotrophic and biotrophic pathogens [41]. Therefore, it is possible that certain NLPs have a role that is independent of cytotoxicity. One such plausible role can be attributed to NLPs by its structural similarity to lectins [42]. Lectins are defined by their ability to bind various carbohydrates such as β-1-3-glucans that are present in the cell wall of filamentous pathogens. Therefore, NLPs being structurally similar to lectins, may bind to cell wall glucans and thereby suppress the recognition of the pathogen by the host surveillance systems. However, this possibility needs further investigation.

Both the AbrNLPs contained bonafide signal peptides and were demonstrated to be secretory in nature using a yeast secretion trap assay. However, their subcellular localisation in *N. benthamiana* was contrasting. While AbrNLP1 was associated with the plasma membrane and the apoplastic space, AbrNLP2 was found not only in the plasma membrane but also in the cytoplasm and nucleus. It is interesting to note that variation in the localisation of NLPs within the same species has not been reported to date. However, different NLPs from different pathogens have been shown to have different subcellular locations. For example, NLPs from *B. cinerea* were associated with the nucleus and nuclear membrane [43]. A type II NLP from *F. oxysporum* was also localised to the cytoplasm using an immunogold-labeling technique [20]. NLPs from the obligate biotroph *Plasmopara viticola* were also found to be localised within the cytoplasm and nucleus as is the case of AbrNLP2 [44]. However, we could not distinguish whether AbrNLP2 is transported into the nucleus or it accumulates at the nuclear membrane (due to its membrane binding affinity).

AbrNLP2 was also shown to induce PTI in both non-host (*N. benthamiana*) and host (*B. juncea*) using the secreted peroxidase assay. The quantum of PTI was higher in the non-host, *N. benthamiana*, which is concordant with the literature that PTI responses against a pathogen are stronger in a nonhost vs. a natural host [45, 46]. Previous studies have demonstrated that NLPs from various fungi, bacteria, and oomycetes act as potent activators of PTI in plants [47]. Further, they have shown that a 24 aa peptide (nlp24) from a conserved region in the NPP1 domain is essential for triggering immune responses [39]. The 24 aa peptide region is highly conserved and is retained in its functional form in AbrNLP1 and AbrNLP2. Recent studies in *Arabidopsis* have shown that RLP23 contributes to resistance against *B. cinerea* via recognition of BcNLP1 and BcNLP2 [48]. However, they also showed that RLP23 is not essential for resistance against *Alternaria brassicicola*, since the NLP from *A. brassicicola* is not expressed at early stages of infection. Therefore, there may be other RLP proteins that may be responsible for the recognition of NLPs from the *Alternaria* species.

In conclusion, we identified two NLPs from *A. brassicae* and functionally characterised their necrosis inducing ability. AbrNLP1 and AbrNLP2 are the first NLPs from the *Alternaria* genus to be functionally characterised. We showed that AbrNLP2 could induce necrosis and PTI effectively in both host and nonhost species. Our work has shown that AbrNLPs are functionally and spatially (localisation) distinct and may play different but important roles during the pathogenesis of *A. brassicae*.

## Supplementary Data

**Supplementary Table S1**: List of Ascomycetes used for the phylogenetic tree construction and their nutrition-based lifestyles

**Supplementary Table S2**: List of primers used in the study and their sequences

**Supplementary Table S3**: List of *Alternaria* species, number of NLPs, and their nutrition-based lifestyle

